# Appropriate data segmentation improves speech encoding models

**DOI:** 10.1101/2024.07.13.603356

**Authors:** Ole Bialas, Edmund C. Lalor

## Abstract

In recent decades, research on the neural processing of speech and language increasingly investigated ongoing responses to continuously presented naturalistic speech, allowing researchers to ask interesting questions about different representations of speech and their relationships. This requires statistical models that can dissect different sources of variance occurring in the processing of naturalistic speech. One commonly used family of models are temporal response functions (TRFs) which can predict neural responses to speech as a weighted combination of different features and points in time. TRFs model the brain as a linear time-invariant (LTI) system whose responses can be characterized by constant transfer functions. This implicitly assumes that the underlying signals are stationary, varying to a fixed degree around a constant mean. However, continuous neural recordings commonly violate this assumption. Here, we use simulations and EEG recordings to investigate how non-stationarities affect TRF models for continuous speech processing. Our results suggest that non-stationarities may impair the performance of TRF models, but that this can be partially remedied by dividing the data into shorter segments that approximate stationarity.

## Introduction

Traditionally, studies on the neural processing of speech and language relied on repetitive presentations of prototypical words or sentences that are carefully manipulated along a single dimension of interest (Hamilton and Huth, 2020). However, while those studies were successful in probing and identifying different modules of speech processing, their artificial and univariate approach makes them incapable of capturing the multi-stream, distributed nature of speech processing (Bouton et al., 2023). One way to overcome this limitation is to record brain responses to continuous narrative stories that present speech in its natural complexity. Analyzing this data requires statistical models that are able to dissect and compare different acoustic, phonetic and semantic representations of speech.

One commonly used family of such models are temporal response functions (TRFs), which estimates the brain’s response to a specific stimulus, or stimulus feature via deconvolution (Crosse et al., 2016). While, in theory, the TRF method works on indefinitely long continuous data, in practice, the data is usually divided into multiple segments. However, there exists no consensus or standard procedure with respect to the duration of those segments. While many studies analyze responses to segments of narrative stories on the order of minutes (Di Liberto, O’sullivan, and Lalor, 2015; Broderick et al., 2018; Gwilliams et al., 2022), others use shorter segments lasting tens of seconds (Golumbic et al., 2013) or even single sentences (Cheung et al., 2016). The way the data is segmented usually depends on the experimental process (e.g., whether the participants listen to individual sentences or long segments of a story) and how this affects the model’s outcome is not considered. This is somewhat negligent because there are several ways in which data segmentation may affect the performance of TRF models.

### Effects of data segmentation

TRFs are linear models that map specific stimulus features to the ongoing brain response. This mapping is estimated by multiplying the covariance matrix for predictor and estimand with the predictor’s autocovariance matrix (see equation 2). When the TRF is estimated across multiple segments, the covariance matrices are averaged across all segments and the TRF is obtained from the average matrices. The validity of this procedure hinges on the assumption that this converges on a stable estimate of the average covariance matrix. However, if there are only few segments, a single outlier may substantially alter the average. If the number of segments is larger, the effect of any single one is reduced. Thus, segmentation regulates overfitting to extreme values.

Segmentation also affects model fitting. Typically, fitting a TRF involves optimizing the regularization parameter λ which penalizes large model weights and prevents overfitting. The optimal value for λ is found by testing multiple candidate values by randomly splitting the data into train and test set, fitting the TRF on the former and evaluating its accuracy by predicting the latter. This requires that both test and train set are representative of the overall trends in the data so that the model can generalize from one set to the other. Representative subsets may be obtained more reliably if many short segments are randomly sampled from the recording.

Of course, segmenting the data into ever shorter bits may impair model accuracy if, at some point, the segments become too short to reliably estimate the covariance matrices. With too little data, the individual covariance matrices might represent the noise in the respective segment, rather than the overall trend in the data. Finally, a key reason for why segment duration matters is that the TRF method treats the brain as a linear time-invariant (LTI) system which implicitly assumes that the modeled signal is a stationary process whose statistical properties are stable across time such that the particular time we observe it is of no relevance (Florescu, 2014).

### Are neural recordings stationary?

Several studies examined the stationarity of EEG recordings by testing whether their amplitude distribution is Gaussian (Campbell et al., 1967; Elul, 1969) or whether observed sequences of values above or below the mean are longer than would be expected by chance (Kawabata, 1976; Cohen and Sances, 1977). They agreed that, above a certain duration, EEG recordings could not be considered stationary, even though the critical duration varied between 2 and 25 seconds (for a review, see Gonen and Tcheslavski, 2012). One study found no difference in the (non-)stationarity between EEG recordings during auditory stimulation, a cognitive task, and resting state, suggesting that these properties are mainly determined by the spontaneous portion of the EEG signal (Kipiński et al., 2011). However, in lack of a ground truth, the origin of non-stationary trends is difficult to assert, and it is of minor importance to the question of appropriate data segmentation.

One can also reason about the stationarity of neural recordings based on the signals’ power spectra. It is generally acknowledged that neural recordings exhibit a 1/f-like power spectrum, meaning that there is an inverse linear relationship between log power and log frequency (He, 2014; Voytek et al., 2015; Meisel et al., 2017). This leads to the contradiction that a 1/f-process can not be stationary, since Rayleigh’s energy theorem states that a signal’s energy is equal to the integral of the power spectrum from negative to positive infinity. For a 1/f-distribution, however, the integral diverges because power approaches infinity as the frequency approaches 0. The alternative is to consider the signal as a non-stationary process where the signal can be integrated over a discrete time and frequency range but a single time-independent energy value can not be obtained (Keshner, 1982; West and Shlesinger, 1990).

In summary, neural time-series exhibit temporal and spectral characteristics of non-stationary processes. This is intuitively plausible, since the brain is a dynamic system where complex networks interact across multiple temporal and spatial scales and the signals recorded from this system mix with noise from various physiological and artificial sources.

### Optimal segmentation

Taken together, the above points suggest that there is an optimal segment duration, where the data can be considered approximately stationary and the effect of outlier segments is suppressed while the individual segments are still long enough to reliably estimate their covariance and autocovariance matrices. Here, we systematically investigate the effect of data segmentation on the performance of TRF models. We use generative simulations to demonstrate that data segmentation improves prediction accuracy in the presence of non-stationary noise. Furthermore, we analyze EEG recordings of neural responses to continuous naturalistic speech and show that segmentation substantially improves prediction accuracy for many participants without negatively affecting the others. We thus recommend that future studies that use TRF models to predict EEG responses to continuous speech use segments of about 10 seconds ^1^.

## Methods & Materials

### Temporal response function

Under our model, the observed neural response is expressed as the weighted sum of the stimulus feature across multiple time lags, defined by the equation:

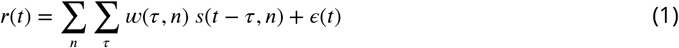

Where *r*(*t*) is the response observed at time *t* and *s*(*t* − *τ, n*) is the stimulus feature at time lag *τ*. The effect of the feature at time lag *τ* is given by the weight *w*(*n, τ*) and the error term *ε*(*t*) contains the residual response unexplained by the model. The vector *w* that contains the weights across all time lags is called the temporal response function (TRF). The TRF is obtained via regularized regression,which, in matrix notation, can be written as:

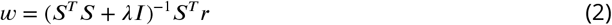

Where (*S*^*T*^ *S*)^−1^ is the inverted autocovariance matrix of the stimulus,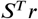 is the covariance matrix of stimulus and response and λ*I* is a diagonal regularization matrix. The value of λ is optimized for model accuracy, defined as the average correlation (Pearson’s r) between the observed response *r*(*t*) and its prediction 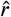(*t*) which is obtained by convolving stimulus features and TRF:

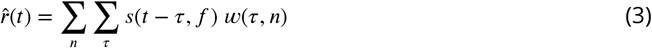

A more detailed mathematical description of the TRF, including multi-variable models, can be found in Crosse et al., 2016.

### Modeling EEG responses to speech

We analyzed data from 19 participants (age 19-38, 13 male) who listened to segments of a classic work of fiction, read by a single American male speaker, which lasted between 170 and 200 seconds. Their brain activity was recorded using a 128-channel ActiveTwo EEG system (BioSemi) at a sampling rate of 512 Hz. All data, which were originally published in two previous studies (Di Liberto, O’sullivan, and Lalor, 2015; Broderick et al., 2018), are available online. We filtered the recordings between 1 and 20 Hz using a non-causal hamming-window band pass filter and downsampled to 64 Hz using MNE-Python (Gramfort et al., 2013). Next, we used the random sample consensus algorithm to identify channels as bad if they were poorly predicted by their neighbors (Bigdely-Shamlo et al., 2015), interpolated them using spherical splines and re-referenced all channels to the global average. We predicted the EEG responses from the spectro-temporal characteristics of the stimulus by band-pass filtering the audio material between 20 and 9000 Hz, dividing it into log-spaced bands (Glasberg and Moore, 1990) and computing the absolute Hilbert envelope for each spectral band, all using the soundlab Python package (Schönwiesner and Bialas, 2021). To test how the number of parameters and the degree of spectral detail affect model accuracy, we represented the stimulus with 1, 8 and 16 spectral bands and repeated the modeling procedure for each representation.

The TRF is fit to the data by testing multiple candidate values for λ and selecting the one that maximizes prediction accuracy. For each of nine log-spaced values between 10^−5^ and 10^3^, the data segments are repeatedly split into train and test set using 5-fold cross validation. For each split, the TRF is estimated on the train set and evaluated by predicting the test set. Model accuracy for a given value of λ is obtained by averaging prediction accuracy across all channels and splits. All modeling was done using the mTRFpy toolbox (Bialas, Dou, and Lalor, 2023).

To estimate the effect of data segmentation on model accuracy, we repeat the above process while dividing the same 50 minutes of data into segments of length 120, 60, 40, 30, 15, 10, 5, 2 and 1 second(s). We excluded the data from further analysis if the accuracy of the best model was below the threshold of 0.01, which affected data from one participant.

### Simulation

We use a simple generative model to test the accuracy of TRF models under different conditions. We define the transfer function for simulating the neural response using a Gabor wavelet, obtained by modulating a sinusoidal with a Gaussian function:

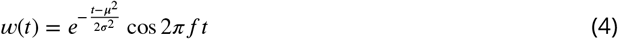

Where *μ* and *σ* are the mean and standard deviation of the Gaussian, of is the sinusoidal’s frequency and *t* is time. The blue curve in Figure 1a shows an example of such a wavelet with *μ* = 0.1 s, *σ* = 0.01 s and *f* = 4 Hz.

**Figure 1.**
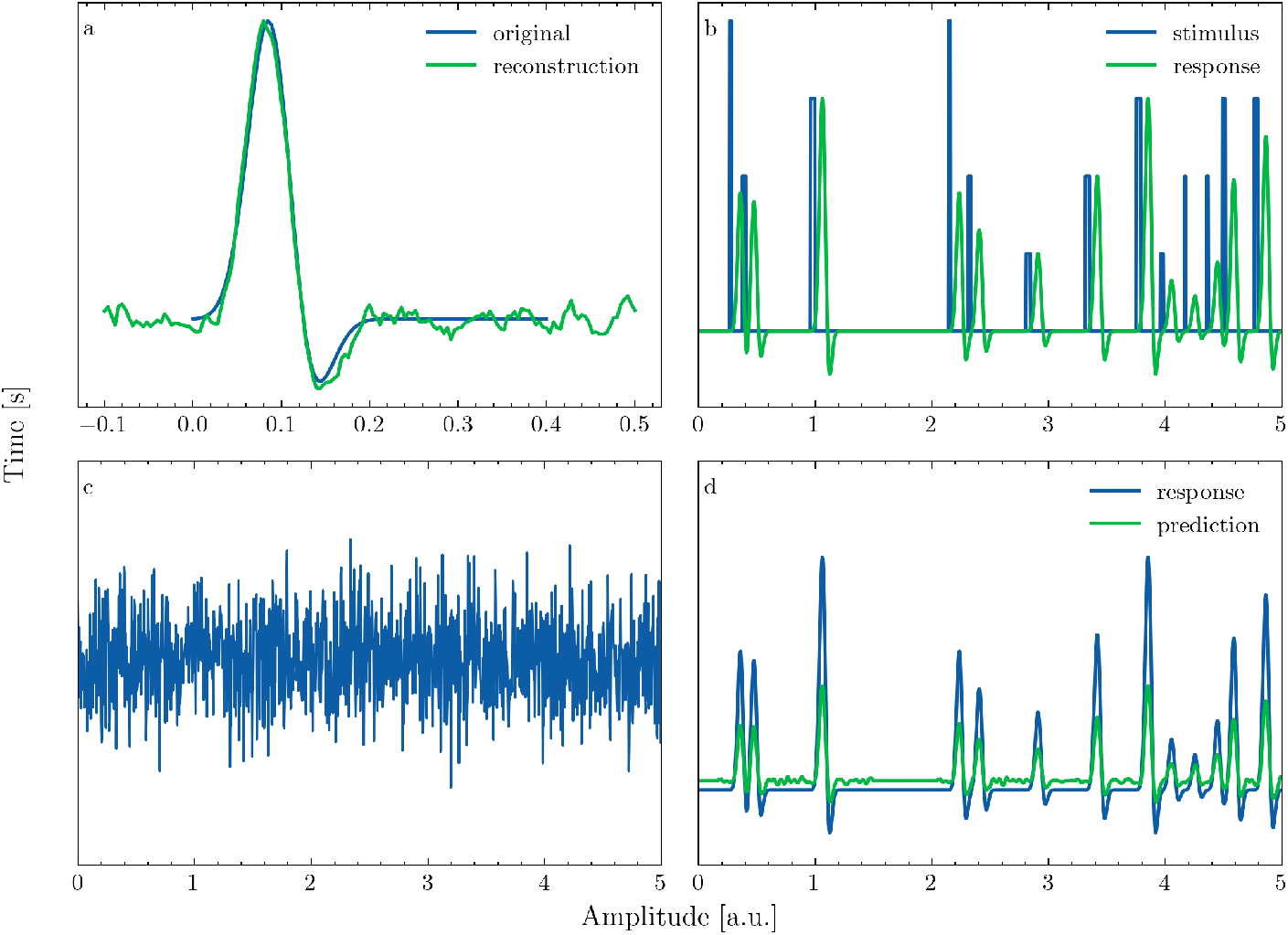
Generative simulation framework. **a**: Original wavelet used for generating and its reconstruction from the noisy response. **b**: Randomly generated stimulus sequence and simulated response obtained by convolving stimulus and wavelet. **c**: Simulated response with Gaussian noise added. **d**: Response predicted using the reconstructed TRF and original response, prior to adding noise.

We simulate a stimulus by generating a sequence of randomly spaced square pulses, where the free parameters are the range of amplitude and width of the pulses (Fig. 1b, blue line). Then, we convolve this stimulus with the wavelet kernel (Fig. 1b, green line) and add noise (Fig. 1c) to obtain the simulated neural response as defined in equation 1. Finally, we fit a TRF to reconstruct the original wavelet (Fig. 1a green line). To estimate the accuracy of the reconstructed TRF, we generate a new stimulus sequence and convolve it with the original and reconstructed TRF and calculate Pearson’s correlation coefficient for the two responses (Fig. 1d).

To determine how violating the assumption of stationarity affects model accuracy, we compare a scenario where the added noise is Gaussian to one where the noise has a 1/f distribution. 1/f noise is generated by computing the Fourier transform of white noise, imposing a 1/f distribution and computing the inverse Fourier transform of the modulated spectrum (for examples of Gaussian and 1/f noise see panels a and b of Figure 2). Note that the simulation represents the best-case scenario where non-stationarity is only introduced by the added background noise. If non-stationary background noise impairs model performance, this problem would certainly be amplified if the generating process itself would vary over time.

**Figure 2.**
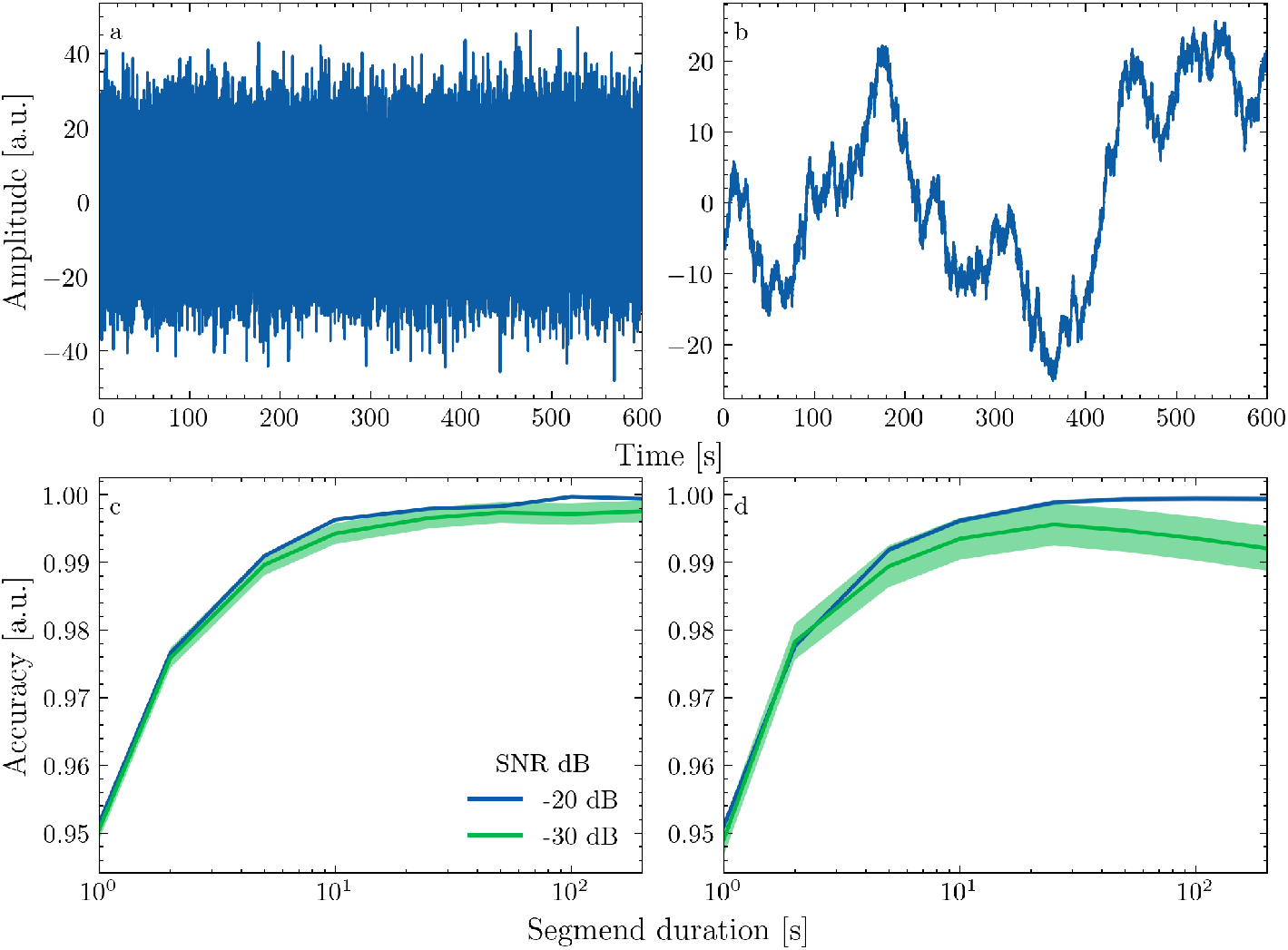
Simulation results: **a** and **b** show samples of Gaussian and 1/f noise. **c** and **d** show normalized reconstruction accuracy as a function of data segment length for two different SNRs.

For both simulations, we divide the data into segments of 200, 100, 50, 25, 10, 5, 2 and 1 second(s), fit a TRF to 1000 seconds of training data and evaluate it on another 1000 seconds of testing data generated from the same model. Since the model is a stochastic process, we repeated the simulation 1000 times.

## Results

The aim of using a generative model to simulate data was to examine the effect of segmentation on model accuracy against a known ground truth. The results show that, when the noise is Gaussian, segmentation into ever shorter bits gradually reduces model accuracy (Fig. 2a). This makes perfect sense since the advantages of segmentation discussed earlier (representative sampling, robustness to outliers, ensuring stationarity), do not apply when dealing with homogeneous data from a stationary process. This trend is independent of the signal-to-noise ratio (SNR), although the model’s variability increases at lower SNRs. However, when the noise is non-stationary, segmentation initially improves prediction accuracy if the SNR is sufficiently low (Fig. 2b). Still, further segmentation decreases prediction accuracy, meaning that there is an optimal length where segments can be considered approximately stationary without substantially reducing model accuracy ^2^.

To test how segmentation affects models trained on actual neural data, we predicted EEG recordings of brain responses to continuous naturalistic speech from the spectrogram of the stimulus while dividing the data into ever shorter segments. To test how the effect of segmentation depends on the model’s flexibility, we repeated the analysis for three different levels of spectral detail. Because prediction accuracy differed strongly between participants and models, we normalized each model’s accuracy, for each participant, by the maximum for that participant, such that the results reflect the relative changes in model accuracy with segment duration. On average, decreasing segment duration from 120s to 10s improved prediction accuracy by roughly 10 to 15 percent on average (Fig. 3a. The improvement was larger for models with more spectral bands (and hence free parameters). Decreasing segment duration below 5s rapidly reduced prediction accuracy. The optimal value of the regularization parameter λ was decreased with segment duration (Fig. 3b). This shows that averaging across segments regularizes the model such that models fit on a larger number of short segments require less additional regularization via λ.

**Figure 3.**
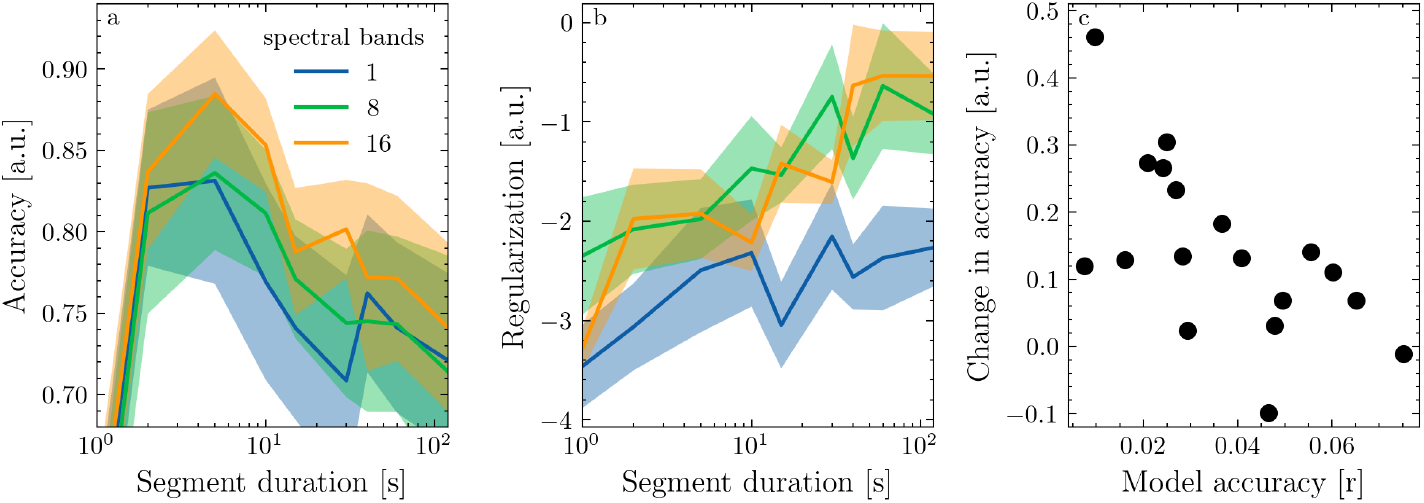
Effect of data segmentation on models for EEG responses to speech. **a**: normalized prediction accuracy as a function of segment duration for three models differing in spectral detail. **b**: optimal (log-transformed) λ decreases with segment duration. **c**: Relative difference in prediction accuracy between models fit on 120s and 10s segments for all participants as a function of their respective prediction accuracy.

The effect of segmentation on model accuracy was highly variable across subjects. To quantify this variability, we compared prediction accuracy between models fitted on 120s and 10s segments for each participant. Models were significantly more accurate when data were segmented into 10 s (r=0.035, SD=0.018) compared to 120s (r=0.031, SD=0.019) segments (two-tailed paired t-test, t=5.0, p=0.0001). What’s more, there was a negative relationship (linear regression, *r* = −0.62, *p* = 0.006) between the relative change in accuracy due to segmentation, and the highest prediction accuracy for each participant (Fig. 3c). This suggests that segmentation was especially effective for participants where model fitting was suboptimal (likely due to non-stationary trends and outliers in the data). Despite the large variability in the effect of segmentation, there was only one participant for whom segmentation slightly reduced model accuracy while, for several participants, it improved prediction accuracy by up to 30 percent.

As we elaborated earlier, one way in which segmentation can affect model accuracy is by reducing the effect of outliers if the number of segments is increased, the effect of any single segment is reduced. To illustrate this point, we selected one of the participants where segmentation had a large effect and estimated the TRF for each segment individually after dividing the data into 120 and 5 (the optimal duration for that participant) second segments, respectively (Fig. 4 a&b). Here, we used only a single spectral band (i.e. the broadband envelope) to avoid arbitrarily selecting features. We identified outliers by sorting the segments by mean absolute TRF weight and selecting the largest five percent. We then computed the TRF for 120 and 5 second segments on the full data and with the outliers removed.

**Figure 4.**
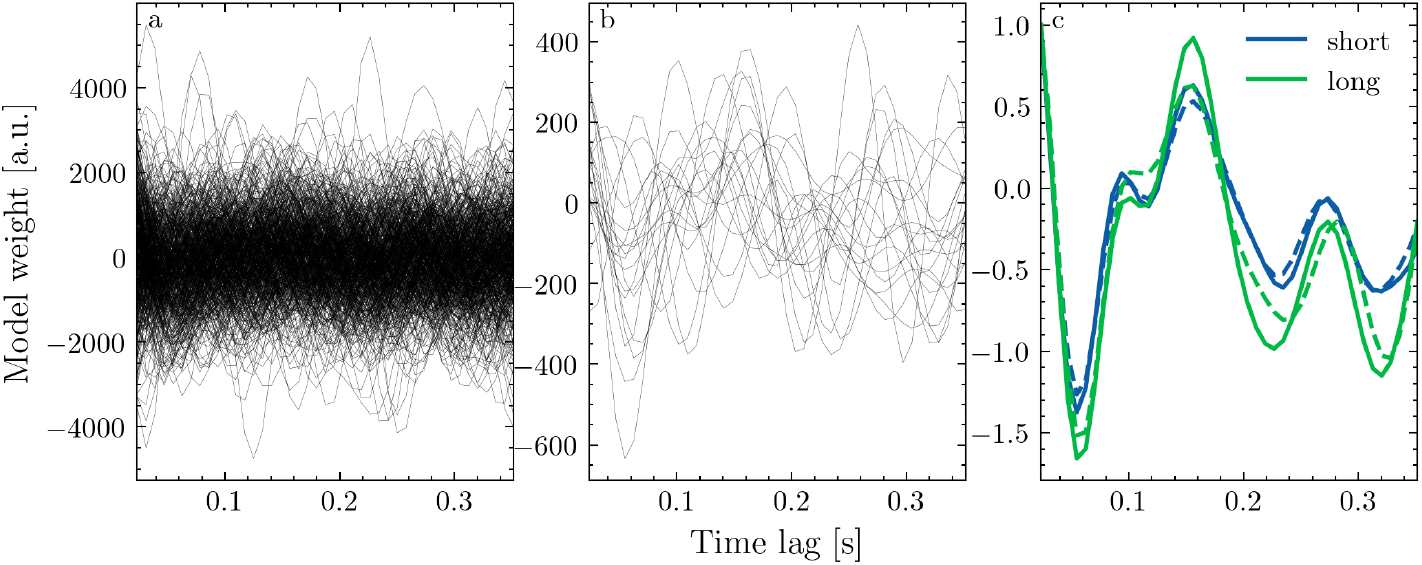
Segmentation regularizes the effect of outliers. **a&b**: TRF for each segment at a fronto-central channel after dividing the data into 5s and 120s segments, respectively. **c**: Average TRF across all segments before (solid) and after (dashed) removing five percent of outliers.

Figure 4c shows the respective TRFs for one fronto-central EEG channel with (solid line) and without (dashed line) outliers. While omitting 5 percent of outlier segments changed the mean absolute weight by 7% for 5 s segments, omitting the same amount of data changed the mean absolute weight by 13 % for 120 s segments. This shows that increasing the number of segments can reduce the effect of outliers on the model’s outcome. Note that this is merely an illustration, the optimal segment duration for isolating outliers will vary across recordings.

## Discussion

We demonstrated that segmentation improves the accuracy of TRF models that predict EEG recordings of brain responses to continuous naturalistic speech. We used a generative model to test the TRFs ability to predict data under different kinds of noise. While dividing the data into ever shorter segments gradually reduced prediction accuracy under stationary noise, segmentation initially improved prediction accuracy when the noise was non-stationary. Analysis of actual EEG recordings revealed a similar trend. This suggests that dividing the data in shorter, standardized, segments remedies the negative effect of non-stationary trends in the data. Optimally, one would choose the longest possible duration where segments can be considered approximately stationary.

For our analysis of brain responses to continuous speech, dividing the data into segments between 5 and 10 seconds yielded the most accurate models. Naturally, one wonders if this generalizes to other EEG data sets. The negative relationship between the effect of segmentation and model accuracy suggests that noise, rather than stimulus related activity, is responsible for nonstationary trends in the data. This is in line with previous findings that the (non-)stationary of EEG recordings does not vary systematically between resting state, auditory stimulation and cognitive tasks (Kipiński et al., 2011). Thus, while the optimal segment duration will certainly depend on the recording conditions (e.g., clinical or mobile recordings may deviate from stationarity more strongly and require more segmentation), it should be independent of the experimental task (or lack thereof). Thus, we cautiously recommend 10 seconds as a default segment duration for encoding models on continuous EEG data but implore researchers to be vigilant about how this may affect their results.

Finally, one may object that dividing the data into segments of equal length is an overly simplistic way of dealing with deviations from stationarity in neural recordings. Maybe, there are extended segments where the recordings are perfectly stationary that would be unnecessarily interrupted by segmentation. There are more sophisticated algorithms that use wavelet transform, fractal dimensions or source separation to adaptively segment neural recordings based on their statistical properties (Anisheh and Hassanpour, 2011; Azami et al., 2015; Haddad et al., 2018). However, implementation of these procedures requires substantial technical knowledge, while dividing the data into normalized segments of equal length can be done in two lines of code and has very little negative consequences.

## Footnotes

1 We don’t necessarily mean to suggest that the experiment needs to be organized in 10 second trials, rather that the data should be restructured prior to model fitting.

2 Note that, since the simulation lacks any physiological plausibility, the absolute values contain no relevant information. For example, the absolute SNRs are very low because a lot of samples in the simulated response are 0, reducing the average power.

